# Evaluating hot water immersion as a quarantine method: impact on *Obama nungara* egg capsule viability

**DOI:** 10.1101/2025.10.14.682045

**Authors:** Johan Watz

## Abstract

Globalization has intensified the spread of invasive species through human-mediated transport, particularly via horticultural trade. The South American land planarian *Obama nungara* has become a widespread invasive soil-dwelling predator in Europe, preying mainly on earthworms and potentially disrupting soil processes. In Sweden, the Environmental Protection Agency recommends heat treatment as a preventive measure against this species. This study tested the hypothesis that a 15-minute 40 °C water bath is lethal to *O. nungara* egg capsules. Thirteen egg capsules were collected from six adult individuals. Every second capsule was subjected to heat treatment, while the remaining served as controls. None of the six heat-treated capsules hatched, whereas five of the seven untreated egg capsules hatched, each produced between five and seven hatchlings after approximately 24 days. These results demonstrate that the recommended heated water bath is effective not only against adults but also against *O. nungara* egg capsules. In addition, the study provides basic ecological information regarding the species’ reproductive biology. Given the species’ invasive potential and its capacity to spread via potted plants, these findings support the implementation of thermal quarantine treatments in nurseries and plant trade to prevent establishment and spread of this invasive flatworm in northern Europe.

## Introduction

Globalization has facilitated the spread of alien species worldwide, driven by human activities such as travel, trade and agroforestry, which have introduced numerous organisms to new regions (Early et al. 2016, Hulme 2021). Introduced species often negatively impact native ecosystems. Soil animals such as earthworms, snails, predatory flatworms and ants have increasingly been recognized as major invasive threats. These organisms can alter nutrient cycling, outcompete or prey upon native fauna and disrupt vegetation. Many invasions occur through human-mediated transport, particularly via potted plants, highlighting the need for effective quarantine measures.

The land planarian *Obama nungara* (Carbayo et al., 2016) is native to South America, primarily the Atlantic Forest of Brazil and Argentina (Boll & Leal-Zanchet 2016, Lago-Barcia et al. 2018). It has invaded several European countries, with introductions largely linked to human activity (Lago-Barcia et al. 2018, Justine et al. 2020, Negrete et al. 2020). Invasive populations have mainly been found in gardens, nurseries, greenhouses and occasionally in natural areas adjacent to human-modified habitats (Justine et al. 2020, Negrete et al. 2020). Genetic analyses indicate multiple independent introductions, often associated with the horticultural trade (Lago-Barcia et al., 2018), especially through transportation of potted plants (Lago-Barcia et al. 2018, Negrete et al. 2020). The effect of local dispersal is likely minor, but may occur by slow crawling under moist conditions (Justine et al. 2020).

*O. nungara* is carnivorous, preying mainly on earthworms but also consuming other soil macrofauna such as gastropods (Boll & Leal-Zanchet 2016, Roy et al. 2022). Gut-content metabarcoding confirms earthworms as the dominant prey in invaded ranges (Roy et al. 2022), but the species is likely a generalist predator, opportunistically using available food resources (Noël et al. 2025b). The species is hermaphroditic and reproduces sexually, laying egg capsules (Lago-Barcia et al. 2018, Justine et al. 2020). Egg capsule production supports rapid local population buildup in favourable habitats (Noël et al. 2025a). Although detailed fecundity rates, generation times and survival under variable climates remain incompletely quantified, populations have been shown to reach high densities in gardens and nurseries (Noël et al. 2025a). Potential ecosystem-level impacts, as a result of earthworm predation, include reduced soil aeration and altered nutrient cycling, although long-term consequences in natural habitats are still uncertain.

The exact upper and lower thermal and desiccation tolerances are not fully quantified. Populations persist in temperate European climates, which indicates some thermal adaptability (Negrete et al. 2020). Hot water treatment has emerged as a promising method to mitigate invasions. Immersion of plants in hot water can kill different soil pests with minor effects on plant health. For example, another invasive flatworm, *Platydemus manokwari* (De Beauchamp, 1963), submerged for 5 min in 43 °C water showed a 100 % mortality rate (Sugiura 2008). Given the significant ecological impacts of invasive soil animals, such quarantine approaches may be crucial for protecting ecosystems and preventing further biological invasions.

The first Nordic observation of *O. nungara* was in Malmö, Sweden, in 2024 (Bouguerche et al. 2025, de Waart et al. 2025), and the Swedish Environmental Protection Agency since recommends heat treatment as a method to prevent its spread. Based on pilot testing on *O. nungara* adults in plant nurseries, the recommendation is to submerge the plant pot in a 40 °C water bath until the entire root ball reaches this temperature and keep it at this temperature for 15 min. This treatment is assumed to kill adult and juveniles (Sugiura 2008). The effect of heat treatment on flatworm egg capsules has not been evaluated, but we hypothesize that heat treatment indeed is effective against *O. nungara* egg capsules. Here, I report results from a laboratory experiment investigating a potential lethal effect of a 15-min-long 40 °C water bath.

## Methods

The experiment was carried out at Karlstad University, laboratory authorized by the Swedish Environmental Protection Agency to conduct research on *O. nungara* (Permit No: SE-IAS 25014). The temperature in the room has held constant at 15 °C. Adult *O. nungara* (Figure 1) were provided by the plant nursery Hallbergs Plantskola (Nossebro, Sweden) and kept individually in 0.6 L plastic containers equipped with pieces of moist paper towels. The *O. nungara* were fed earthworms (*Dendrobaena veneta* Rosa, 1886) *ad libitum*. Terrestrial planarians are known to be able to store sperm and produce egg capsules also in isolation (Ducey et al. 2005; Noël et al., 2015b). Egg capsules used in this study (n = 13; Figure 1) were produced by six adult individuals between 15 August and 19 September 2025 (min = 1, max = 3 egg capsules per individual); found egg capsules were collected directly on sight and their diameters measured to the nearest mm.

**Figure 1.**
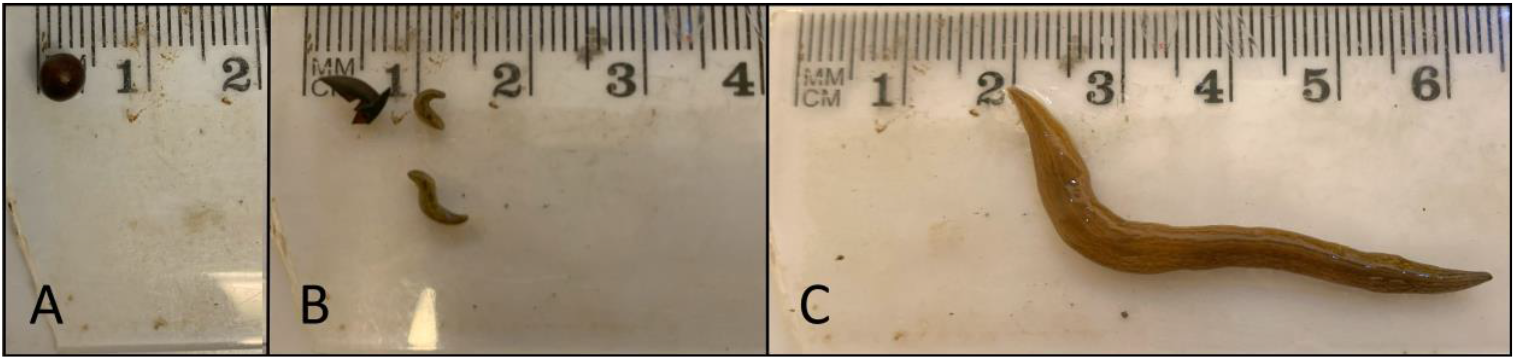
The invasive planarian *Obama nungara* at different life stages: egg capsule (A), newly hatched juveniles (B) and an adult (C).

Every second egg capsule found was submerged for 15 min in a 40 °C water bath directly on discovery, using a small aquarium fish net and a 500 mL glass beaker placed in a heating cabinet. Either after heat treatment or directly on discovery (controls), egg capsules were placed in separate 50 mL Faclon tubes. The tubes were provided with a piece of moist paper towel and punctured lids to allow air into the tube. The tubes were kept in a 14 L holding box with non-tight-fitting black plastic bag around the box to minimize light input. The egg capsules were checked on average every second day, and when hatchlings appeared (Figure 1) their number and the day were recorded. The experiment was terminated on 14 October, observing intact, non-hatched egg capsules for at least 36 days. Non-hatched egg capsules at this time were assumed not to be viable. For egg capsules that hatched, we calculated incubation time (time from the observation of a new egg capsule to its hatching) and evaluated the effect of heat treatment on viability using a 2 × 2 Fisher’s exact test.

## Results

The egg capsule diameter ranged from 3 to 5 mm (mean ± SD = 4.2 ± 0.7 mm). Out of the seven untreated egg capsules, five capsules hatched, each producing between five and seven hatchlings (mean ± SD = 5.8 ± 0.8). It took between 21 and 25 days from the discovery of a new egg capsule and its hatching (mean ± SD = 23.6 ± 1.7 days). None of the six heat-treated egg capsules hatched (one-tailed Fisher’s exact test, n = 13, p = 0.01), effectively demonstrating that the heat treatment was lethal. Most heat-treated egg capsules looked the same as the untreated ones during incubation, except one heat-treated egg capsule that burst after 13 days, releasing a whitish sludge.

## Discussion

The results presented here support the hypothesis that heat treatment is effective also against *O. nungara* egg capsules. In addition, the study provides basic ecological data regarding the species’ life history, such as egg capsule incubation time (∼ 24 days at 15 °C), number of hatchlings per egg capsule (5–6) and a rough estimate of interval between egg capsule laying (4 – 13 days). In a recent study on *O. nungara* by Noël et al. (2025b), the most fecund individual laid 10 egg capsules in one month (∼ 1 every 4 days), a similar frequency as reported here. Moreover, Noël et al. (2025b) observed more hatchlings per egg capsule (mean = 9, min – max = 7 – 12) and the incubation time in their study was shorter (2 – 3 weeks) than reported in this study. They used a temperature regime with 18 °C during the day and 14°C at night, therefore the average temperature was 1 °C higher in their study, which may partly explain the discrepancies. Differences in prey food source may also play a role, and different haplotypes connected to different invasion occasions (Justine et al. 2020, Bouguerche et al. 2025) also potentially results in variation in some life history parameters.

*O. nungara* is spreading rapidly over Europe and has potentially not yet reached its northern distribution limit (Fourcade 2021), and a scenario with rapid and extensive spread in the Nordic countries is possible (*sensu* the case of the Spanish slug *Arion vulgaris* Moquin-Tandon, 1855; Hatteland et al. 2013), with detrimental effects on the native fauna and soil-related ecosystem services. *O. nungara* was found in a park Malmö, Sweden, in November 2024 (Bouguerche et al., 2025), and was subjected to emergency measures by the Swedish government (under Section 11, Ordinance 2018:1939 on Invasive Alien Species). This meant that the plant trade was required to carry out measures targeting the species. To prevent further spread, effective quarantine measures targeting its vectors need to be developed and evaluated. The present study provides evidence that a 15-min-long 40 °C water bath, recommended by the Swedish Environmental Protection Agency, effectively kills off potential egg capsules hidden in root balls of potted plants.

## Acknowledgements

Many thanks to Maria Nielsen at Hallbergs Plantskola organizing the collection of *O. nungara* for the experiment. Nya Eriksbo Plantskola, Splendor Plant, Stångby Plantskola, Lackalänga Trädgård generously provided *O. nungara*. Thank you! Karlstad University, Department of Environmental and Life Sciences, funded the study.

## References

Bouguerche, C., Roth, J., & Justine, J. L. (2025). Flatworms in the land of the midnight sun: first record of the invasive species Obama nungara (Platyhelminthes, Geoplanidae) in Sweden, its northernmost location in continental Europe. bioRxiv preprint, 2025.03.24.644988. 10.1101/2025.03.24.644988

de Waart, S. A., Vanhove, M. P. M., Justine, J. L., & Kmentová, N. (2025). Going Dutch: European distribution of non-native land flatworm species belonging to Geoplaninae and Bipaliinae with focus on the Netherlands. NeoBiota, 99: 285–321. 10.3897/neobiota.99.145703

Ducey, P. K., West, L. J., Shaw, G., & De Lisle, J. (2005). Reproductive ecology and evolution in the invasive terrestrial planarian Bipalium adventitium across North America. Pedobiologia, 49(4), 367–377. 10.1016/j.pedobi.2005.04.002

Early, R., Bradley, B. A., Dukes, J. S., Lawler, J. J., Olden, J. D., & Blumenthal, D. M. (2016). Global threats from invasive alien species in the twenty-first century and national response capacities. Nature Communications, 7, 12485. 10.1038/ncomms12485

Fourcade, Y. (2021). Fine-tuning niche models matters in invasion ecology. A lesson from the land planarian Obama nungara. Ecological Modelling, 457, 109686. 10.1016/j.ecolmodel.2021.109686

Hatteland, B. A., Roth, S., Andersen, A., Kaasa, K., Støa, B., & Solhøy, T. (2013). Distribution and spread of the invasive slug Arion vulgaris Moquin-Tandon in Norway. Fauna norvegica, 32, 13–26. 10.5324/fn.v32i0.1473

Hulme, P. E. (2021). Unwelcome exchange: International trade as a direct and indirect driver of biological invasions worldwide. One Earth, 4(5), 616–629. 10.1016/j.oneear.2021.04.015

Lago-Barcia, D., Fernández-Álvarez, F. Á., Brusa, F., Rojo, I., Damborenea, C., Negrete, L., Grande, C., & Noreña, C. (2018). Reconstructing routes of invasion of Obama nungara (Platyhelminthes: Tricladida) in the Iberian Peninsula. Biological Invasions, 21(2), 289–302. 10.1007/s10530-018-1834-9

Justine, J.-L., Winsor, L., Gey, D., Gros, P., & Thévenot, J. (2020). Obama chez moi! The invasion of metropolitan France by the land planarian Obama nungara (Platyhelminthes, Geoplanidae). PeerJ, 8, e8385. 10.7717/peerj.8385

Justine, J.-L., Gastineau, R., Gey, D., Robinson, D. G., Bertone, M. A., & Winsor, L. (2024). A new species of alien land flatworm in the Southern United States. PeerJ, 12, e17904. 10.7717/peerj.17904

Negrete, L., Lenguas Francavilla, M., Damborenea, C., & Brusa, F. (2020). Trying to take over the world: Potential distribution of Obama nungara (Platyhelminthes: Geoplanidae), the neotropical land planarian that has reached Europe. Global Change Biology, 26(8), 4623–4635. 10.1111/gcb.15208

Boll, P. K. & Leal-Zanchet, A. M. (2016). Preference for different prey allows the coexistence of several land planarians in areas of the Atlantic Forest. Zoology, 119(3), 162–168. 10.1016/j.zool.2016.04.002

Noël, S., Fourcade, Y., Roy, V., Bonnet, G., & Dupont, L. (2025a). Population dynamics of the exotic flatworm Obama nungara in an invaded garden. Ecology and Evolution, 15(1), 1–11. 10.1002/ece3.70827

Noël, S., Fourcade, Y., Roy, V., Gigon, A., & Dupont, L. (2025b). Experimental assessment of the predation pressure by the exotic flatworm Obama nungara in its introduced range. Current Zoology, 71(2), 1–11. 10.1093/cz/zoaf025

Roy, V., Ventura, M., Fourcade, Y., Justine, J.-L., Gigon, A., & Dupont, L. (2022). Gut content metabarcoding and citizen science reveal the earthworm prey of the exotic terrestrial flatworm, Obama nungara. European Journal of Soil Biology, 108, 1–9. 10.1016/j.ejsobi.2022.103449

Sugiura, S. (2008). Hot water tolerance of soil animals: Utility of hot water immersion in preventing invasions of alien soil animals. Applied Entomology and Zoology, 43(2), 207–212. 10.1303/aez.2008.207

